# The Glutamatergic Projection from the Substantia Nigra Pars Reticulata to the Dorsal Raphe Nucleus Facilitates Social Hierarchy in Mice

**DOI:** 10.1101/2024.11.12.623319

**Authors:** Yanzhu Fan, Shaoxiang Ge, Lidi Lu, Wenjun Niu, Zhiyue Wang, Xiaoguo Jiao, Guangzhan Fang

**Affiliations:** Chengdu Institute of Biology, Chinese Academy of Sciences. No.23 Qunxian Nan Street, Tianfu new area, 610213, Chengdu, Sichuan, China; School of Life Sciences, Hubei University, No. 368 Youyi Avenue, 630062, Wuhan, Hubei, China

**Keywords:** social hierarchy, substantia nigra pars reticulata, dorsal raphe nucleus, neural circuit, glutamatergic neurons

## Abstract

Social hierarchy constitutes a fundamental organizational characteristic among various social species, significantly influencing individual survival, health, and reproductive success within these societies. Neurons in the substantia nigra pars reticulata (SNr) exhibit extensive connectivity with the dorsal raphe nucleus (DRN), a critical structure implicated in social interaction, reward processing, and the establishment of social rank. However, the specific neuronal types within the SNr, as well as the associated neural circuits that regulate social dominance, remain inadequately characterized. This study aims to elucidate the crucial role of SNr glutamatergic neurons in the establishment and maintenance of social hierarchy in mice. Employing fiber photometry, we observed that the activation of SNr glutamatergic neurons increased during the initiation of effortful behaviors in the tube test. Further investigations revealed that optogenetic activation or chemogenetic inhibition of the glutamatergic neural circuit connecting the SNr to the DRN induced upward or downward shifts in social ranks, respectively. Additionally, our findings indicate that the activation of SNr glutamatergic terminals in the DRN reduces anxiety levels in mice. Collectively, these results underscore the critical role of the glutamatergic pathway from the SNr to the DRN in regulating social hierarchy. This work enhances our understanding of the functions of SNr glutamatergic neurons in both physiological contexts and neurological disorders.

## Introduction

The social hierarchy serves as a fundamental organizing principle in many animal societies, including insects, fish, reptiles, rodents, and primates(Byrne and Bates, 2010; Grosenick et al., 2007; Paz-y-Miño et al., 2004). This organizational framework delineates individual rankings within a group, established through recurrent social interactions such as aggressive encounters or behavioral displays(Stagkourakis et al., 2018; Tibbetts et al., 2022; Wang et al., 2014). An individual’s position within this hierarchy is frequently associated with various fitness-related outcomes. For example, social dominance status significantly influences an individual’s health, survival, and reproductive success(Buston, 2003; Dugatkin and Druen, 2004; Fan et al., 2023; Nagy et al., 2010; Piper, 1997; Sapolsky, 2005; Smith, 1981; Zhou et al., 2018). Moreover, all group members benefit from the stability provided by dominance hierarchies, which mitigate agonistic behaviors and, in turn, conserve energy and reduce the risk of physical injury(Chou et al., 2016; Dewsbury, 1982; Drews, 1993; Milewski et al., 2022; Sapolsky, 2005; Zink et al., 2008). Consequently, elucidating the central neural mechanisms underlying social hierarchies is of paramount importance for comprehending both individual life histories and population dynamics. Despite this significance, the processes through which dominance hierarchies are established and maintained remain inadequately understood.

Numerous brain regions have been implicated in the regulation of social dominance, including the prefrontal cortex (PFC), amygdala, hippocampus, hypothalamus, striatum, and lateral habenula(Chou et al., 2016; Dwortz et al., 2022; Padilla-Coreano et al., 2022; Tan et al., 2018; Wang et al., 2014; Watanabe and Yamamoto, 2015). Notably, the PFC has been identified as the principal regulatory center for social hierarchy, processing information regarding social status received from upstream brain regions and modulating the expression of dominant behaviors in downstream areas(Padilla-Coreano et al., 2022; Wang et al., 2011; Zhang et al., 2022; Zhou et al., 2017). Among the identified neural circuits relevant to social dominance, the reward- and motivation-related mesolimbic dopamine system is particularly prominent(Ghosal et al., 2019). In murine models, social submission appears to coincide with a reduction in motivational drive, suggesting that social dominance is intricately linked to motivational processes(Kunkel and Wang, 2018). Importantly, motivational and reward-related processes are largely regulated by the mesolimbic system, leading to the inference that this system may play a critical role in the formation of social hierarchies. Consistent with this notion, various subcortical structures, including the substantia nigra pars reticulata (SNr), dorsal raphe nucleus (DRN), and amygdala, are involved in the representation of social rank signals in some species(Dwortz et al., 2022). For instance, dopamine (DA) signaling is elevated in response to alterations in status signals within the SNr of green anole lizards (*Anolis carolinensis*)(Korzan et al., 2006), indicating a potential role for the SNr in the establishment and maintenance of dominance hierarchies.

The SNr functions as a pivotal hub for the integration of diverse upstream input signals, coordinating multifaceted functions such as motor control, cognition, habit formation and reward(Antal et al., 2014; Arber and Costa, 2022; Dudman and Krakauer, 2016; Galaj et al., 2020; Jin and Costa, 2010; Kravitz et al., 2010; Liu et al., 2020; Partanen and Achim, 2022; Rizzi and Tan, 2019; Schmidt et al., 2013; Sobczak et al., 2021; Wicker et al., 2019; Yasuda and Hikosaka, 2015; Zhou and Lee, 2011). While the majority of neurons in the SNr are GABAergic, the population also includes glutamatergic and dopaminergic neurons, as well as those utilizing both glutamate (Glu) and dopamine as neurotransmitters. In comparison to the well-characterized GABAergic neurons of the SNr, the role of glutamatergic neurons remains less elucidated; however, some SNr glutamatergic neurons may share developmental and functional relationships with their GABAergic counterparts. The DRN, a critical component of the mesolimbic system, has been identified as the primary source of serotonin (5-HT) in the brain(Ren et al., 2018; Sengupta et al., 2017; Sengupta and Holmes, 2019). Previous studies have established a robust association between the serotoninergic system and social rewards, as well as the expression of social hierarchies across a variety of species, including crustaceans, reptiles, rodents, and primates(Cases et al., 1995; Coccaro, 1992; Edwards and Kravitz, 1997; Korzan and Summers, 2004; Larson and Summers, 2001; Li et al., 2016; Raleigh et al., 1991; Sandi and Haller, 2015; Terranova et al., 2016; Watanabe and Yamamoto, 2015; Yeh et al., 1996). For instance, the injection of 5-HT into the hemolymph of both lobsters and crayfish reduces the likelihood of retreat and extends the duration of aggressive interactions(Huber et al., 1997; Livingstone et al., 1980). Moreover, experimentally enhancing serotonergic activity has been shown to facilitate the acquisition of dominance in monkeys, whereas reductions in this activity correspondingly diminish dominance(Raleigh et al., 1991). Similar effects have been documented in humans, where the administration of 5-HT influences social dominance(Moskowitz et al., 2001). Collectively, these findings underscore the potential role of the DRN and the serotonergic system in the regulation of social dominance. Notably, both serotonergic and GABAergic neurons within the DRN receive glutamatergic inputs from the SNr(Dorocic et al., 2014; Xu et al., 2021). Thus, we hypothesize that the glutamatergic pathway extending from the SNr to the DRN (SNr ^Glu^-DRN) may play a significant role in regulating social hierarchy.

To investigate this hypothesis, we employed fiber photometry to measure the activity patterns of SNr^Glu^ neurons in conjunction with specific behaviors exhibited by mice in the competitive tube test. The results indicated heightened activity levels of these neurons during instances of effortful behavior in the tube test. Furthermore, we utilized cell-type-specific optogenetic and chemogenetic methodologies to manipulate SNr^Glu^ neurons and evaluate their role in social competition. Our findings demonstrate that optogenetic activation or chemogenetic inhibition of the SNr ^Glu^-DRN pathway induces an increase or decrease in social ranks, respectively. Collectively, these findings highlight the pivotal role of the glutamatergic pathway from the SNr to the DRN in the regulation of social hierarchy.

## Results

### Population activity of SNr^Glu^ neurons increased during effortful behavioral epochs in social contest scenarios

To investigate the real-time population activity of SNr^Glu^ neurons during social competition in the tube test, we bilaterally administered rAAV-CaMKIIα-GCaMP6f, which encodes the fluorescent calcium indicator GCaMP6f, or rAAV-CaMKIIα-EYFP into the SNr of mice (Fig. 1a, b). Subsequently, we bilaterally implanted optical fibers to facilitate the recording of calcium signals from SNr^Glu^ neurons, while concurrently utilizing a camera to monitor the animals’ behavior (Fig. 1c). By capturing calcium signals from a randomly selected unilateral SNr, we observed a marked increase in SNr^Glu^ neuronal activity immediately upon the initiation of pushing and counter-pushing during direct confrontations with an opponent in the tube (Fig. 1d). In contrast, no significant alterations in calcium signals were detected when mice traversed the tube without any opponent or when EYFP-expressing mice were subjected to the tube test (Fig. 1e, f). The observed Ca^2+^ dynamics implying that SNr^Glu^ neurons are dynamically activated during effortful behavioral epochs, such as push-initiation and push-back, and that their population activity is correlated with active social contests. These findings indicate the relationship between SNr^Glu^ neuronal activity and social competition and status.

**Figure 1.**
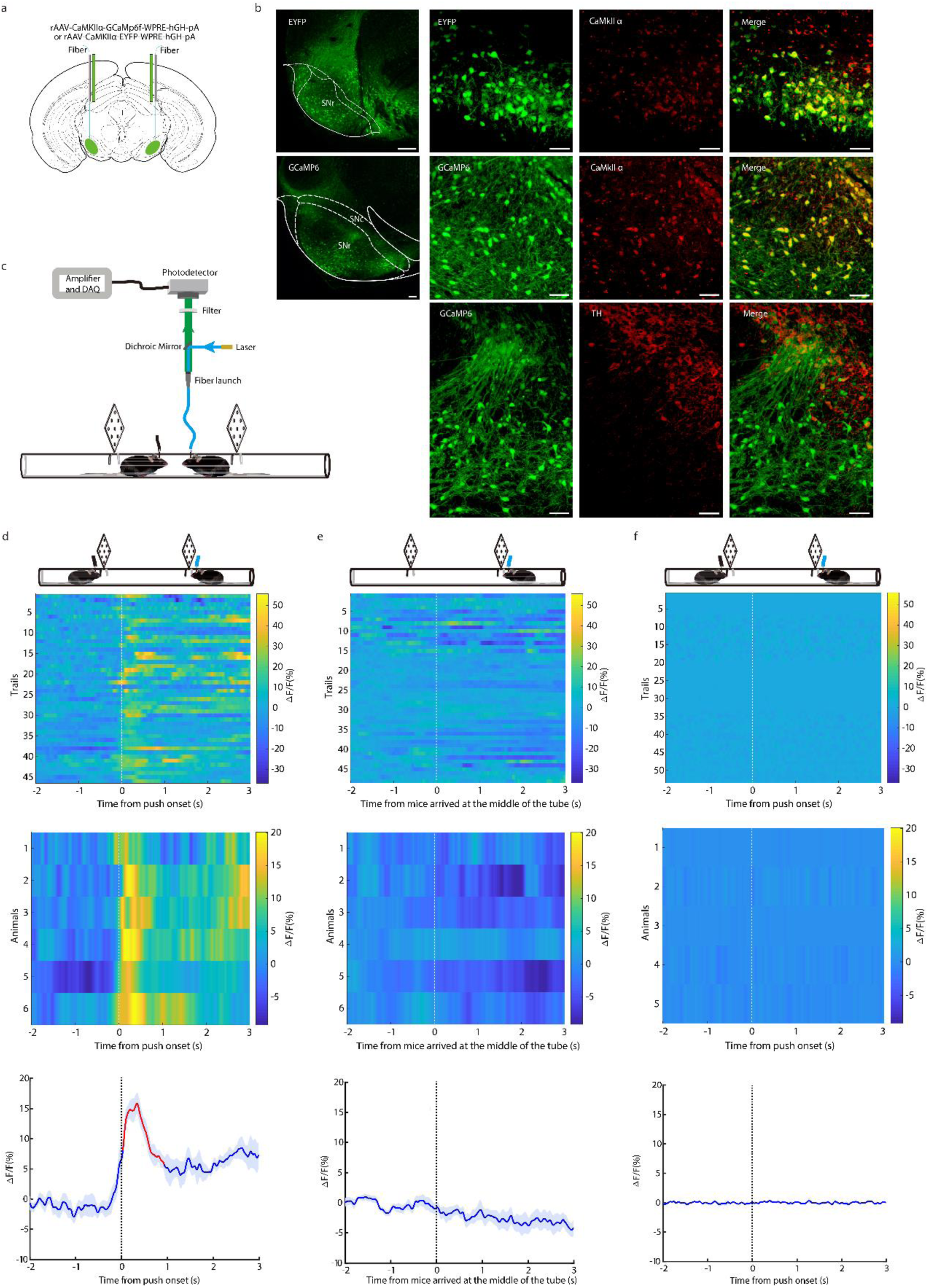
Ca²⁺ activity of SNr^Glu^ neurons during push-initiation and push-back in the tube test. **a**. Schematic representation of the CaMKIIα-GCaMP6f viral construct, injection site, and optic fiber placement over the SNr. **b**. Left, a representative coronal section of the SNr shows the expression of EYFP or GCaMP6f. Scale bar, 200 μm. Right, immunostaining confirming the specificity of EYFP and GCaMp6f expression in glutamatergic neurons (not dopaminergic neurons). Scale bar, 40 μm. **c**. Schematic of the fiber photometry setup used to record SNr^Glu^ neuron calcium activity during the tube test. **d**. Ca²⁺ signals aligned with the initiation of pushes when mice expressing GCaMP6f encounter their cagemates. The upper panel shows a heatmap of Ca²⁺ signals aligned to push-initiation and push-back, each row represents one trial (n = 46 trials from 6 mice). The middle panel displays a heatmap representing the average Ca²⁺ signals for each animal across all trials (n = 6). The lower panel shows mean Ca²⁺ transients associated with pushes for the entire test group (n = 6), with solid lines indicating mean and shaded areas indicating SEM. A red segment denotes a statistically significant increase from baseline (*p* < 0.05; permutation test). **e**. Ca²⁺ signals aligned with the time point when mice reached the middle of the tube without an opponent. The upper panel presents data from all trials (n = 48), the middle panel from all animals (n = 6), and the lower panel shows the mean ± SEM of the average Ca²⁺ signals. **f**. Ca²⁺ signals aligned with the initiation of pushes in mice expressing EYFP encountering their cagemates. The upper panel presents data from all trials (n = 53), the middle panel from all animals (n = 5), and the lower panel shows the mean ± SEM of the average Ca²⁺ signals. Abbreviations: SNc, substantia nigra pars compacta; SNr, substantia nigra pars reticulata; TH, tyrosine hydroxylase.

### Optogenetic activation of SNr^Glu^ neurons sufficiently enhances social status in the tube test

To investigate the behavioral responses elicited by the activation of SNr^Glu^ neurons, we bilaterally delivered rAAV-CaMKIIα-hChR2(H134R)-EYFP or rAAV-CaMKIIα-EYFP constructs to the SNr of mice (hereafter referred to as ChR2-EYFP mice and EYFP mice, respectively; see Fig. 2a, b). Subsequently, we implanted optic fibers above the injection sites. Following a minimum of four weeks post-viral injection, we stimulated the randomly selected unilateral SNr in vivo using pulses of 473 nm blue light (20 ms duration, 20 Hz frequency, and 10-20 mW intensity) prior to the mice entering the tube. Photostimulation of the unilateral SNr significantly influenced behavioral performance in the tube test (Fig. 2d-h), resulting in elevated tube test rankings for ChR2-EYFP mice (Fig. 2i-k), whereas no such effects were observed in EYFP mice (Supplementary Fig. 1). In ChR2-EYFP mice, photostimulation markedly enhanced effortful behaviors, including push-initiation (Fig. 2d) and push-back (Fig. 2e), while concurrently reducing the incidence of retreats (Fig. 2f). Notably, photostimulation did not affect stillness (Fig. 2g) or the proportion of time spent resisting (Fig. 2h). Compared to controls, the significantly elevated social ranks were maintained for at least four days (Fig. 2i-k). Collectively, these findings suggest that optogenetic activation of SNr^Glu^ neurons in mice promotes an increase in social hierarchy.

**Figure 2.**
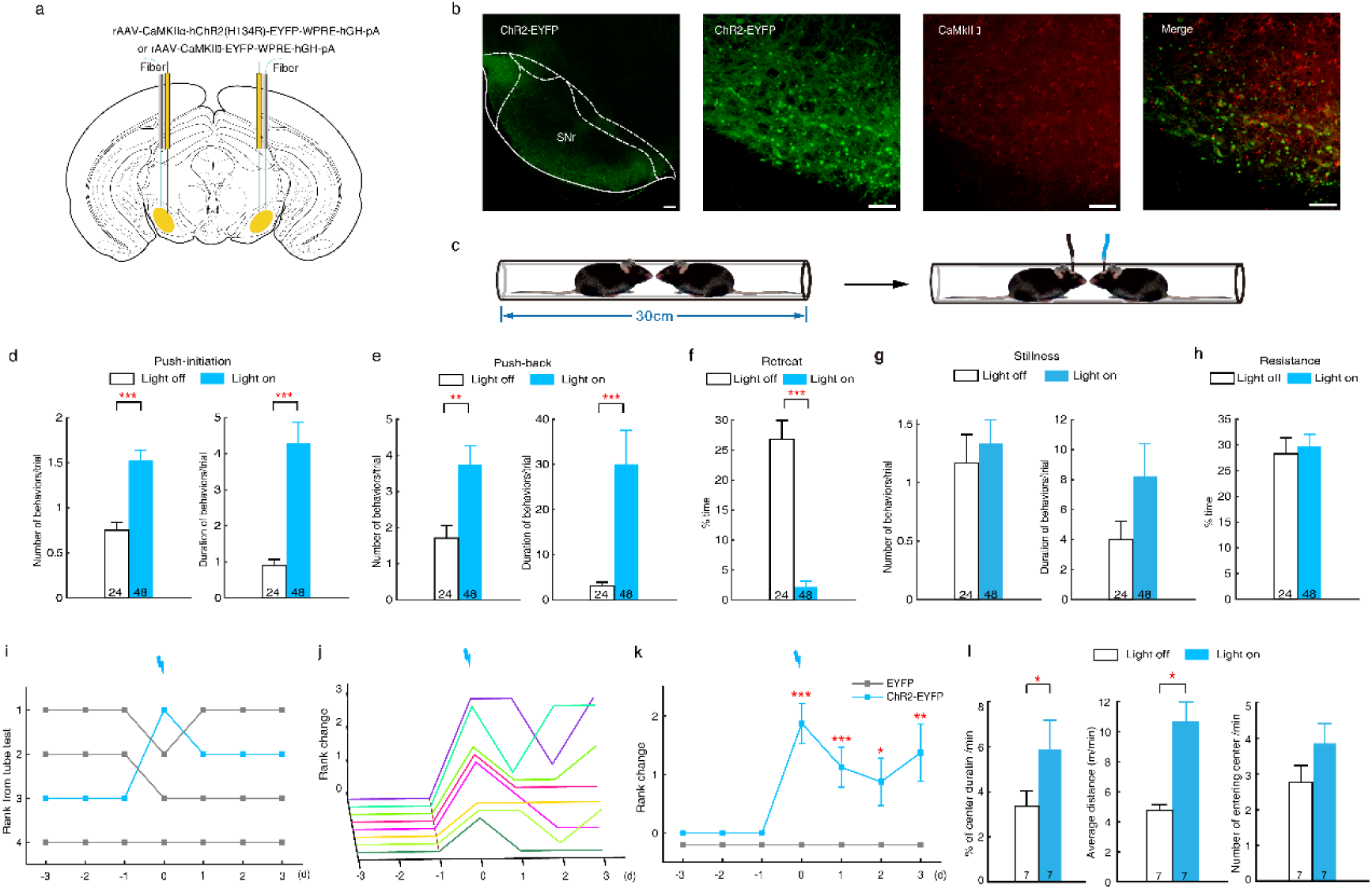
Optogenetic activation of SNr^Glu^ neurons induces winning in the tube test. **a** A schematic representation illustrating the ChR2-EYFP and EYFP viral constructs, injection sites, and optic fiber positioning above the SNr. **b** Left, a representative coronal section of the SNr shows the expression of ChR2-EYFP. Scale bar, 200 μm. Right, immunostaining confirming the specificity of ChR2-EYFP expression in glutamatergic neurons. Scale bar, 40 μm. **c** Diagrams illustrating the tube test with and without optogenetic activation. **d** Quantification of push-initiation occurrences and durations for each trial in the tube test (n=24 for light off and n=48 for light on, Mann-Whitney U test; number: U=267, *p*<0.001; duration: U=155.5, *p*<0.001). **e** Analysis of push-back occurrences and durations for each trial in the tube test (n=24 vs. 48, Mann-Whitney U test; number: U=331.5, *p*<0.01; duration: U=170.5, *p*<0.001). **f** Percentage of time spent retreating (n=24 vs. 48, Mann-Whitney U test; U=39, *p*<0.001). **g** Quantification of stillness occurrences and durations for each trial in the tube test (n=24 vs. 48, Mann-Whitney U test; number: U=546, *p*=0.709; duration: U=501, *p*=0.362). **h** Percentage of time spent resisting (n=24 vs. 48, Mann-Whitney U test; U=546, *p*=0.720). **i** An example demonstrating how the third-ranked mouse in a given cage achieved first place following photostimulation, assessed daily over seven days. **j** Summary of rank changes in ChR2-EYFP mice pre- and post-optogenetic activation of SNr^Glu^ neurons, with each line representing an individual animal. **k** Difference in average rank change between ChR2-EYFP and EYFP mice after optogenetic activation of SNr^Glu^ neurons (n=8 for ChR2-EYFP mice and n=9 for EYFP mice, Mann-Whitney U test; Day 0: U=0, *p*<0.001; Day 1: U=4.5, *p*<0.001; Day 2: U=13.5, *p*=0.007; Day 3: U=9, *p*=0.002). **l** Optogenetic activation of SNr^Glu^ neurons increased center time and locomotion in the open-field test (center time: n=7, paired two-sided t-test, t_6_=-2.95, *p*=0.032; average distance: n=7, Wilcoxon signed rank test, Z_6_=-2.366, *p*=0.018), but did not affect the number of entries into the central area (n=7, paired two-sided t-test, t_6_=-1.887, *p*=0.108). * *p* < 0.05; ** *p* < 0.01; *** *p* < 0.001.

Subsequently, we conducted a series of behavioral assays to further explore the association between various behavioral performances and the optogenetic activation of SNr^Glu^ neurons in mice. During light-on periods, both the time spent in the central area (center time) and the locomotion of mice in the open field test (OFT) significantly increased (Fig. 2l). In contrast, the number of entries into the central area did not exhibit a significant change compared to light-off conditions. These findings suggest that the activation of SNr^Glu^ neurons is sufficient to enhance exploratory and locomotor behaviors. Notably, the duration of time spent in the open arms prior to light activation in the elevated plus-maze (EPM) test was significantly greater than that observed during and after light-on periods; however, no differences in open arm time were found between ChR2-EYFP and EYFP mice (Supplementary Fig. 2a), indicating that optogenetic activation of SNr^Glu^ neurons does not influence anxiety-like behaviors. Furthermore, SNr-photostimulated mice exhibited a typical preference for novel mice in the social memory test (Supplementary Fig. 2b) and displayed normal levels of both aggressive and non-aggressive behaviors toward novel mice in the resident-intruder test (Supplementary Fig. 2c). Additionally, SNr photoactivation did not impact muscle strength (Supplementary Fig.2d). These results collectively indicate that the alterations in social hierarchy observed in mice following the optogenetic activation of SNr^Glu^ neurons are not attributable to changes in social preference, aggressive levels, or forelimb grip strength.

### Chemogenetic inhibition of SNr^Glu^ neurons results in defeat in the tube test

To further validate the critical role of SNr^Glu^ neurons within the competitive context of the tube test, we employed the inhibitory DREADD (designer receptors exclusively activated by designer drugs) system. Specifically, we bilaterally administered rAAV-CaMKIIα-hM4D(Gi)-EGFP or rAAV-CaMKIIα-EYFP into the SNr of mice, subsequently designated as hM4Di-EGFP mice and EYFP mice, respectively (Fig. 3a, b). Following this intervention, Clozapine-N-oxide (CNO, 10 mg/kg) was administered intraperitoneally to each mouse in the experimental group, while their cage mates received an equivalent volume of saline. The tube test was conducted at intervals of 1-1.5, 3-5, 6-8, 24, 48, and 72 hours post-injection. The inhibition of SNr^Glu^ neurons resulted in significant behavioral alterations and a reduction in tube test rankings among the hM4Di-EGFP mice treated with CNO, in comparison to the EYFP mice receiving CNO (Fig. 3c-k) and the hM4Di-EGFP mice receiving saline (Supplementary Fig. 3). At 3.5 hours post-injection, both the number and duration of push initiations (Fig. 3c) and the duration of immobility (Fig. 3e) were significantly diminished in the hM4Di-EGFP mice, while the frequency of push-backs (Fig. 3d), resistance (Fig. 3f), and retreats (Fig. 3g) increased. The significantly reduced rankings compared to controls were sustained for approximately eight hours; however, by 24 hours post-CNO administration, most mice reverted to their original rank positions (Fig. 3i-k).

**Figure 3.**
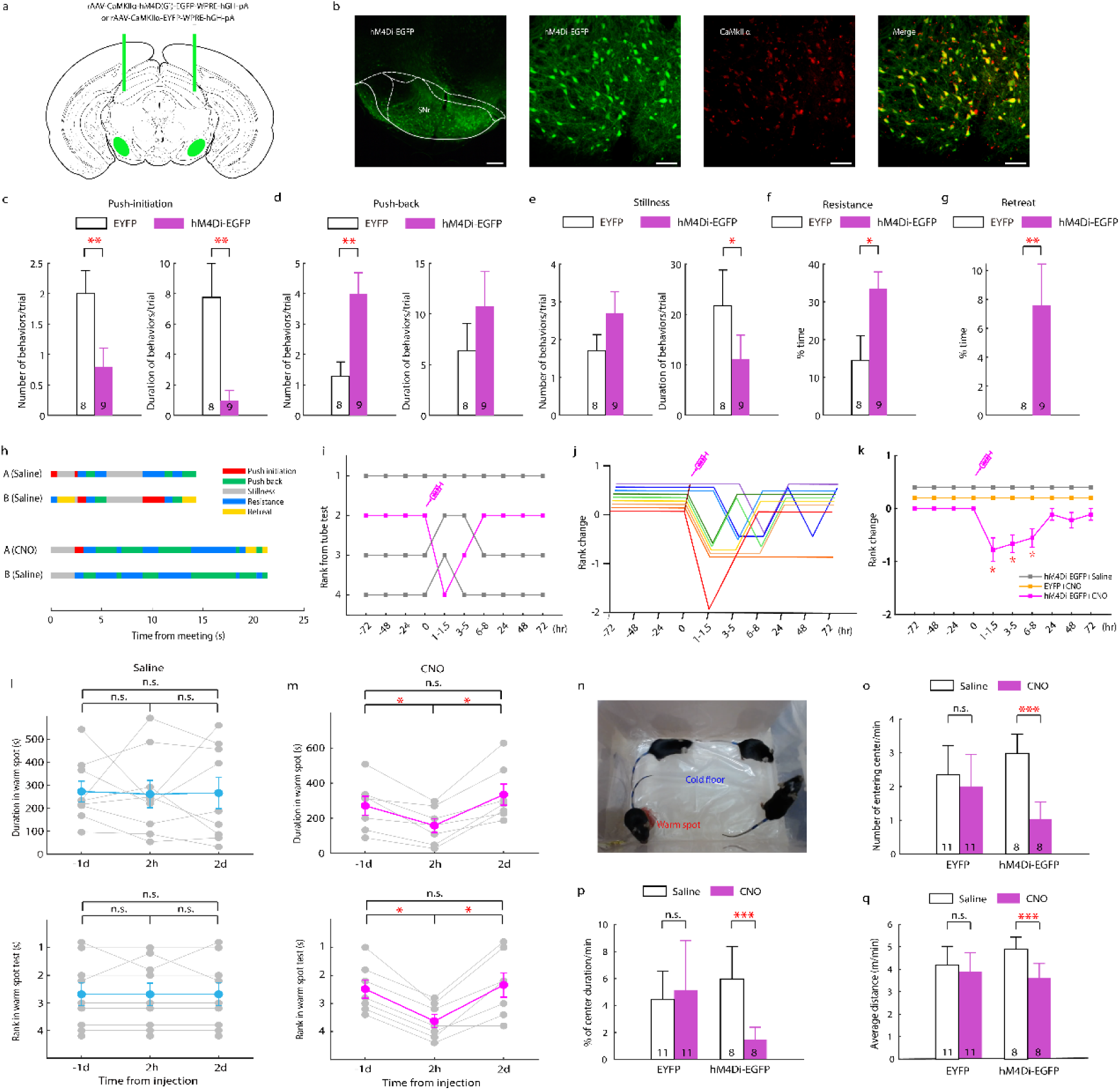
Chemogenetic inhibition of SNr^Glu^ neurons results in defeat in the tube test. **a** A schematic representation of the hM4Di-EGFP and EYFP viral construct and their injection sites in the SNr. **b** Left, a representative coronal section of the SNr, illustrating hM4Di-EGFP expression. Scale bar, 200 μm. Right, immunostaining confirming the specificity of hM4Di-EGFP expression in glutamatergic neurons. Scale bar, 40 μm. **c** Number and duration of push-initiation for each trial in the tube test (n=8 mice for EYFP and n=9 for hM4Di-EGFP, Mann-Whitney U test; number: U=7, *p*=0.004; duration: U=7, *p*=0.005). **d** Number and duration of push-back for each trial in the tube test (n=8 vs. 9, Mann-Whitney U test; number: U=9, *p*=0.008; duration: U=30, *p*=0.563). **e** Number and duration of stillness for each trial in the tube test (n=8 vs. 9, Mann-Whitney U test; number: U=31, *p*=0.621; duration: U=15, *p*=0.043). **f** Percentage of time spent resisting (n=8 vs. 9, Mann-Whitney U test; U=11, *p*=0.016). **g** Percentage of time spent retreating (n=8 vs. 9, Mann-Whitney U test; U=8, *p*=0.003). **h** Behavioral annotation of two tube test trials between the same pair of hM4Di-EGFP mice, pre- and post-CNO injection, with a one-week interval between trials. **i** Tube test outcomes for a cohort of hM4Di-EGFP mice before and after CNO administration to the rank-2 mouse at time 0. **j** Summary of rank changes in hM4Di-EGFP mice following CNO administration to each individual at time 0, with each line representing an individual mouse. **k** Mean change in rank prior to and following injection of CNO or saline (n=9 for hM4Di+CNO, n=7 for hM4Di+Saline while n=8 for EYFP+CNO, Wilcoxon signed rank test, at 1-1.5 hr: Z=-2.333, *p*=0.020; at 3-5 hr: Z=-2.449, *p*=0.014; at 6-8 hr: Z=-2.236, *p*=0.025). **l** and **m** Duration (top) and rank (bottom) in the warm spot test for hM4Di-EGFP mice at 1 day pre-injection, 2 hours post-injection, and 2 days post-injection of saline (duration: n=9, one-way repeated measures ANOVA with Bonferroni correction, F_2,16_=0.023; *p*=0.978; rank: Friedman test, t_2_=0.667; *p*=0.717) or CNO (duration: n=7, F_2,12_=11.902; *p*=0.001; rank: t_2_=11.043; *p*=0.004). **n** Depiction of the warm spot test, where four mice from the same cage compete for a warm corner in a cage with an ice-cold floor. **o-q** Chemogenetic inhibition of SNr^Glu^ neurons led to a reduction in entries into the center, time spent in the center, and locomotion in the open field test for hM4Di-EGFP mice (one-way repeated measures ANOVA with Bonferroni correction; number of entries: n=8, F_1,7_=171.726, *p*<0.001; center time: n=8, F_1,7_=61.080, *p*<0.001; average distance: n=8, F_1,7_=43.053, *p*<0.001) but not for EYFP mice (n=11, *p*>0.05). * *p* < 0.05; ** *p* < 0.01; *** *p* < 0.001.

To explore whether the alterations in tube test rankings induced by chemogenetic inhibition of SNr^Glu^ neurons can be generalized to rankings assessed through other paradigms, we conducted the warm spot test (Fig. 3n). Our findings demonstrate that inhibition of SNr^Glu^ neurons led to a significant reduction in the duration of time spent in the warm spot by hM4Di-EGFP mice at 2 hours post-CNO injection, compared to saline controls (Fig. 3l-m). In contrast, EYFP mice exhibited no significant changes in warm spot occupancy or social hierarchy following either CNO or saline injections (Supplementary Fig. 4). These results indicate that social hierarchical states can be generalized across various social contexts in mice. Taken together, our data suggest that chemogenetic inhibition of SNr^Glu^ neurons contributes to a decline in social dominance, thereby underscoring the essential role of these neurons in the maintenance of social hierarchy.

To assess the potential alterations in exploratory and locomotion behaviors subsequent to the chemogenetic inhibition of SNr^Glu^ neurons, we conducted an open field test on mice. Our results indicate a significant reduction in the number of entries into the center, time spent in the center, and overall locomotor activity among hM4Di-EGFP mice two hours post-CNO injection, in contrast to the saline injection (Fig. 3o-q). Notably, no significant differences in these locomotor parameters were observed in EYFP mice following either CNO or saline injections (Fig. 3o-q). These findings underscore the critical role of SNr^Glu^ neuron activation in sustaining heightened exploratory and locomotor behaviors.

### SNr^Glu^ neurons modulate social hierarchy via the SNr^Glu^-DRN pathway

Next, we investigated the specific output pathways of the SNr, focusing on the glutamatergic neurons. By injecting rAAV-CaMKIIα-EYFP virus into the SNr of wild-type mice, we selectively labeled SNr glutamatergic neurons and their axons with EYFP (Fig. 4a). EYFP-expressing SNr glutamatergic axons were observed in the forebrain and brainstem regions, including the caudate putamen (CPu), ventral posteromedial thalamic nucleus (VPM), zona incerta (ZID), and DRN (Fig. 4b). To assess whether the specific stimulation of SNr glutamatergic projections can modulate social hierarchy, we stimulated the terminals of SNr^Glu^ neurons in these regions in ChR2-EYFP mice, using the protocol from the tube test. Optogenetic activation of SNr^Glu^-CPu,-VPM and -ZID pathways did not affect the social hierarchy of mice (Supplementary Fig. 5). However, activation of the SNr^Glu^-DRN pathway increased effortful behaviors and enhanced social ranking in ChR2-EYFP mice (Fig. 4c-h), consistent with the effects observed upon photostimulation of SNr^Glu^ neurons in the SNr, while no similar effects were observed in EYFP control mice (Supplementary Fig. 6). In ChR2-EYFP mice, photostimulation of SNr^Glu^ terminals in the DRN significantly increased effortful behaviors, including push-initiation (Fig. 4c) and push-back (Fig. 4d), while concurrently decreasing the frequency of retreats (Fig. 4e). However, this photostimulation did not significantly impact stillness (Supplementary Fig. 7a) or resistance (Supplementary Fig. 7b). Notably, the observed elevation in social status was sustained for at least three days in comparison to the control group (Fig. 4g, h). Collectively, these findings indicate that optogenetic activation of the SNr^Glu^-DRN pathway is sufficient to enhance social dominance.

**Figure 4.**
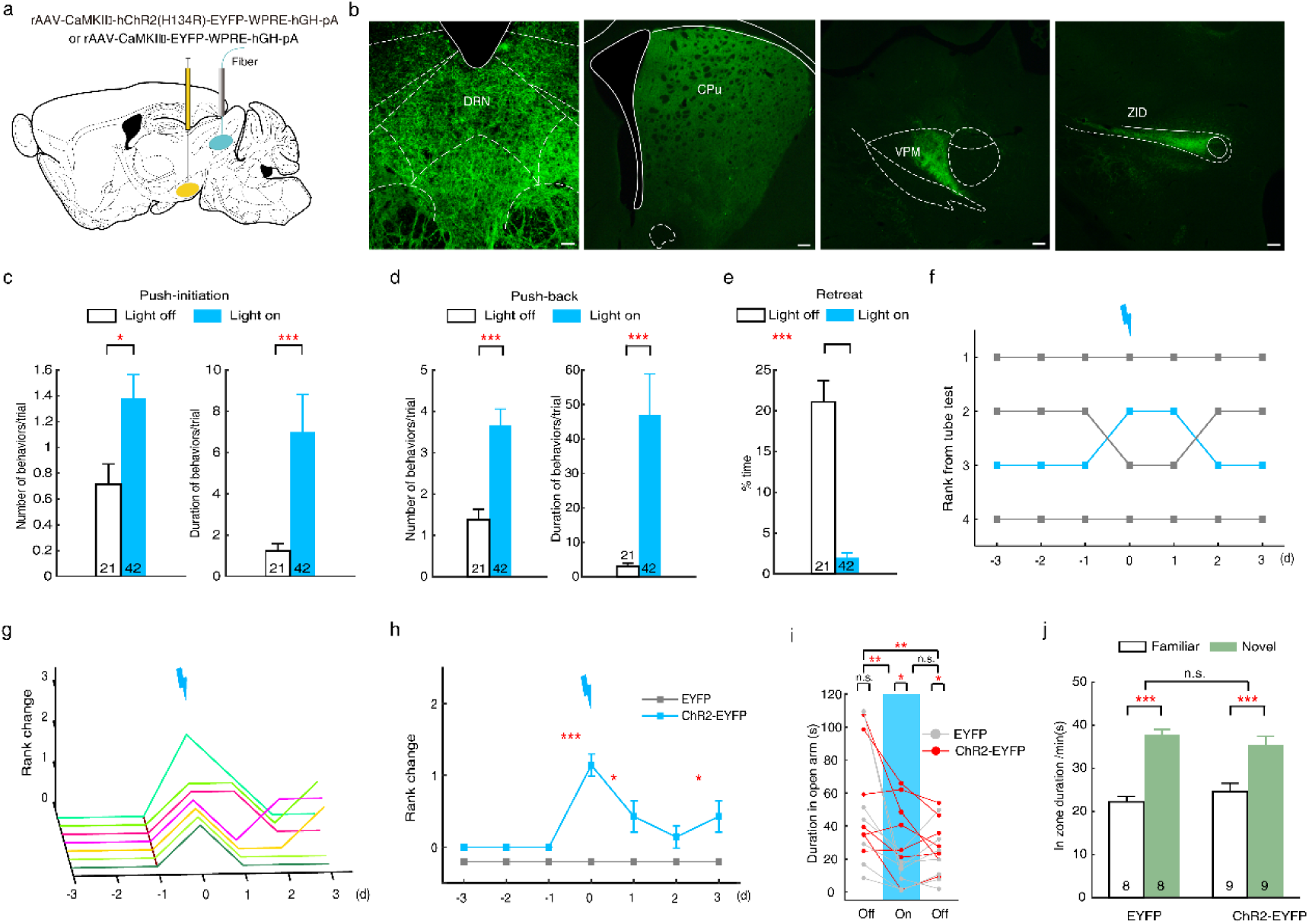
Optogenetic activation of SNr^Glu^-DRN pathway induces winning in the tube test. **a** A schematic representation of ChR2-EYFP and EYFP viral constructs, detailing their injection sites within the SNr, alongside the optic fiber placement above the DRN. **b** Representative SNr terminal sites expressing EYFP. Scale bar, 200 μm. **c** Number and duration of push-initiation for each trial in the tube test (n=21 for light off and n=42 for light on, Mann-Whitney U test; number: U=291, *p*=0.020; duration: U=217.5, *p*=0.001). **d** Number and duration of push-back for each trial in the tube test (n=21 vs. 42, Mann-Whitney U test; number: U=162.5, *p*<0.001; duration: U=92, *p*<0.001). **e** Percentage of time spent retreating (n=21 vs. 42, Mann-Whitney U test; U=23, *p*<0.001). **f** Example of rank position dynamics over a 7-day period within a cohort of mice, illustrating the ascension of the third-ranked mouse to second position following photostimulation of SNr^Glu^ neuron terminals in the DRN. **g** Summary of rank changes in ChR2-EYFP mice pre- and post-optogenetic activation of the SNr^Glu^-DRN pathway. Each trajectory represents a single animal. **h** Comparison of average rank changes between ChR2-EYFP and EYFP mice post-optogenetic activation of the SNr^Glu^-DRN pathway (n=7 for the ChR2-EYFP mice and n=8 for the EYFP mice, Mann-Whitney U test; Day 0: U=0, *p*<0.001; Day 1: U=16, *p*=0.046; Day 2: U=24, *p*=0.285; Day 3: U=16, *p*=0.046). **i** In the elevated plus maze test, the time spent in the open arms decreased over time (n=7 for the ChR2-EYFP mice and 7 for EYFP mice, two-way repeated measures ANOVA with Bonferroni correction; F_2,24_=15.257, *p*<0.001), with ChR2-EYFP mice spending more time in the open arms compared to EYFP controls (F_1,12_=6.610, *p*=0.024). **j** Both ChR2-EYFP and EYFP mice demonstrated a significant preference for novel mice during the social memory test under light-on conditions (n=9 for ChR2-EYFP mice and 8 for EYFP mice, two-way repeated measures ANOVA with Bonferroni correction; F_1,15_=28.896, *p*<0.001), with no significant difference observed between the groups (F_1,15_=0.052, *p*=0.822). Abbreviations: CPu, caudate putamen (striatum); DRN, dorsal raphe nucleus; VPM, ventral posteromedial thalamic nucleus; ZID, zona incerta, dorsal part. \**p* < 0.05; \*\**p* < 0.01; \*\*\**p* < 0.001.

To elucidate the behavioral implications associated with optogenetic activation of the SNr^Glu^-DRN pathway in mice, we next conducted a series of behavioral assays. Notably, we observed that the duration spent in the open arm prior to light activation during the EPM test was significantly greater than that observed during and after light activation (Fig. 4i). In comparison to EYFP mice, photostimulation of the terminals of SNr^Glu^ neurons in the DRN resulted in increased duration in the open arm for ChR2-EYFP mice (Fig. 4i), suggesting that optogenetic activation of the SNr^Glu^-DRN pathway may modulate anxiety levels in mice. Furthermore, photostimulation of this pathway preserved typical preferences for novelty in the social memory test (Fig. 4j), and exhibited a standard range of aggressive and non-aggressive behaviors towards novel conspecifics in the resident-intruder test (Supplementary Fig. 7c). Additionally, metrics such as the number of entries into the central area, time spent in the central area, and locomotion in the OFT under light exposure did not differ significantly compared to the light-off condition (Supplementary Fig. 7d). Importantly, photostimulation of the SNr^Glu^-DRN pathway did not influence forelimb muscle strength (Supplementary Fig. 7e). Collectively, these findings suggest that the SNr^Glu^-DRN circuit may play a critical role in the regulation of both social hierarchy and anxiety levels, thereby indicating a potential link between alterations in social hierarchy and anxiety in mice.

## Discussion

As the largest output nucleus of the basal ganglia, the SNr integrates a multitude of upstream input signals and projects to various brainstem regions, functioning as a central hub for the signal integration and coordination of these targets’ functions(Antal et al., 2014; Zhou and Lee, 2011). Anatomically, the SNr is organized into parallel modules that influence both ascending and descending targets implicated in the regulation of behaviors such as motor control, cognition, habit formation, and reward processing(Arber and Costa, 2022; Dudman and Krakauer, 2016; Galaj et al., 2020; Jin and Costa, 2010; Kravitz et al., 2010; Liu et al., 2020; Partanen and Achim, 2022; Rizzi and Tan, 2019; Schmidt et al., 2013; Sobczak et al., 2021; Wicker et al., 2019; Yasuda and Hikosaka, 2015). Crucially, the SNr occupies a strategic position in the processing of reward-related information, which guides motor responses to secure future rewards(Bryden et al., 2011; Galaj et al., 2020). Given that social interactions related to dominance hierarchies represent one aspect of a broader spectrum of reward-related behaviors, it is plausible that the activities of SNr neurons are associated with social interactions. Consistent with this notion, our findings reveal that the activation and inhibition of SNr^Glu^ neurons exert opposing effects on competitive outcomes. Notably, the heightened activity of SNr^Glu^ neurons during effortful behaviors underscore their critical role in mediating social hierarchy, further reinforcing the essential functions of the SNr in various physiological regulation processes(Antal et al., 2014; Zhou and Lee, 2011). Consequently, our study elucidates a causal relationship between social dominance and SNr^Glu^ neurons, highlighting the fundamental role of the SNr in both social dominance and effortful behaviors.

The current results indicate that activation of the SNr^Glu^-DRN pathway enhances hierarchical dominance in mice, consistent with previous studies highlighting the role of the DRN in social interactions, reward processing, cognition, and aggressive behaviors(Audero et al., 2013; Nautiyal et al., 2015). The serotoninergic system is closed associated with social rewards and the expression of social hierarchies across various species, including crustaceans, reptiles, rodents, and primates(Cases et al., 1995; Coccaro, 1992; Edwards and Kravitz, 1997; Korzan and Summers, 2004; Larson and Summers, 2001; Li et al., 2016; Raleigh et al., 1991; Sandi and Haller, 2015; Terranova et al., 2016; Watanabe and Yamamoto, 2015; Yeh et al., 1996). For instance, the injection of 5-HT into the hemolymph of lobsters and crayfish reduces the likelihood of retreat and prolongs aggressive encounters(Huber et al., 1997; Livingstone et al., 1980). Similarly, dominant male vervet monkeys (*Cercopithecus aethiops*) exhibit approximately double the concentration of 5-HT compared to subordinate individuals(Raleigh et al., 1984). Furthermore, experimentally enhancing serotonergic activity facilitates the acquisition of dominance in monkeys, while reductions in this activity diminish dominance(Raleigh et al., 1991). Comparable effects have been documented in humans, where administration of 5-HT similarly impacts social dominance(Moskowitz et al., 2001). Collectively, these studies underscore the significance of the serotonergic system in regulating social dominance and status. The DRN serves as the primary source of 5-HT in the brain, with serotonergic neurons constituting two-thirds of the total neuronal population and establishing extensive forebrain connections(Ren et al., 2018; Sengupta et al., 2017; Sengupta and Holmes, 2019). Recent investigations have identified that the SNr provides glutamatergic input to both serotonergic and GABAergic neurons within the DRN(Dorocic et al., 2014; Xu et al., 2021). In this context, our study demonstrates that the SNr^Glu^-DRN pathway is involved in the regulating hierarchy dominance. However, further research is necessary to elucidate the specific types of downstream neurons and the synaptic connections implicated in this function.

Hierarchical position significantly influences physiological responses, behavior, and mental health, thereby affecting an individual’s overall emotional state(Bartolomucci et al., 2001; Yin et al., 2023). Specifically, social hierarchy plays a crucial role in determining susceptibility to stress and is recognized as a critical risk factor for the development of psychiatric disorders(LeClair and Russo, 2021; Sandi and Haller, 2015). In numerous species, a U-shaped relationship exists between stress hormone levels and social rank(Dwortz et al., 2022). While dominant individuals may enjoy privileged access to resources, they also bear the burden of maintaining their status, which can lead to increased vigilance and stress(Ferrari et al., 1998; Larrieu et al., 2017). Conversely, subordinate individuals often face challenges associated with restricted access to resources and limited social interactions, potentially inducing anxiety due to uncertainty or perceived threats(Blanchard et al., 1984; Horii et al., 2017). This dynamic could lead to an elevated risk of chronic social defeat stress(LeClair and Russo, 2021) and stress-related psychiatric conditions such as depression and anxiety(Larrieu et al., 2017). Social dominance can be attained through competitive interactions, yet both stress and anxiety can influence the formation of social hierarchies, underscoring their critical roles in determining competitive outcomes among animals(Sapolsky, 2005). Recent studies in both humans(Goette et al., 2015) and rodents(Hollis et al., 2015; van der Kooij et al., 2018) have identified trait anxiety as a crucial predictor of social competitiveness. In humans, individuals of lower social status experience a disproportionately high prevalence of mental disorder(Gilbert and Allan, 1998; Wilkinson, 1999), and exhibit increased mortality rates compared to their higher-ranking counterparts(Ghosal et al., 2019; Sapolsky, 2005). In rat models, highly anxious individuals typically lose territorial contests against low-anxious conspecifics(Horii et al., 2017), while low-anxious individuals are more likely to achieve dominance when cohabitating for extended periods(Davis et al., 2009). Consistent with these observations, the current findings indicate that the activation of glutamatergic projections from the SNr to the DRN not only enhances social status in mice but also alleviates anxiety levels. This suggests a mutually reinforcing relationship between low-anxious status and the acquisition of dominance. Collectively, these findings highlight the SNr^Glu^-DRN pathway as a pivotal neural circuit for modulating anxiety and social dominance, potentially providing insight into the intricate relationship between social hierarchy and anxiety levels(Fan et al., 2023). This relationship may be mediated through distinct neural circuits or receptor mechanisms, emphasizing the need for further research to elucidate the circuitry and synaptic mechanisms underpinning the intrinsic connections between social hierarchy and anxiety levels in mice.

In summary, our study indicates that the activation of SNr glutamatergic neurons is both necessary and sufficient for the rapid induction of winning in social competitions. Notably, this effect is not mediated by an enhancement of aggression or physical strength; rather, it stems from the initiation and maintenance of effortful behaviors. These findings carry significant implications for future research on social behavior and could inform the development of therapeutic interventions for neuropsychiatric disorders, including anxiety. Future studies employing more refined manipulations of SNr glutamatergic neurons and their downstream targets are anticipated to yield a deeper understanding of the neural circuitry that regulates social hierarchy and anxiety.

## Methods

### Animals

In the present study, adult male C57BL/6 mice (aged 8-12 weeks) were obtained from Chengdu Dossy Experimental Animals Co., Ltd. (Chengdu, China). The mice were housed under controlled experimental conditions, featuring a 12-hour light/dark cycle, a temperature maintained at 22 ± 1 °C, and a humidity level of 65 ± 5%, with ad libitum access to food and water. All experimental procedures were conducted in compliance with the guidelines established by the Animal Care and Use Committee of Chengdu Institute of Biology, Chinese Academy of Sciences (permit number: CIBDWLL2020001). The mice were randomly assigned to experimental and control groups. Although a specific statistical method was not employed to predetermine the sample size, the sizes utilized were consistent with those conventionally accepted in the field.

### Surgery

Mice were anesthetized using 1-2% isoflurane in oxygen and positioned in a stereotaxic apparatus (51603, Stoelting, USA) equipped with a heating pad. Anesthesia depth was consistently maintained at a stable level, indicated by the absence of reflexes. Viral vector microinjection was conducted with a microliter syringe (NRS75 RN 5.0 μL (33/20/3), Hamilton, Switzerland) attached to a syringe pump (Legato 130, KD Scientific, USA) securely mounted to the stereotaxic apparatus. A volume ranging from 200 to 500 nl of AAV solution was administered into the SNr (AP = −3.3 mm, ML = ± 1.5 mm, DV = −4.5 mm). Following injection, the micropipette was held in place for 10 minutes before being gently retracted. We used the following high titer AAVs (in genomic particles/ml): AAV2/9-CaMKIIα-GCaMP6f (2.94 × 10^12^, 400 nl/site for optical fiber-based Ca^2+^ recordings), AAV2/9-CaMKIIα-hChR2(H134R)-EYFP (2.86 × 10^12^; 300 nl/site for optogenetic activation), AAV2/9-CaMKIIα-hM4D(Gi)-EGFP (4.89 × 10^12^; 300 nl/site for chemogenetic inhibiton) and AAV2/9-CaMKIIα-EYFP (2.73 × 10^12^; 200 nl/site for anterograde tracing and control groups). Two weeks post-viral injection, two fiber optic cannulas (1.25 mm in diameter; Fiblaser, Shanghai, China) were implanted above the SNr (AP = −3.3 mm, ML = ± 1.5 mm, DV = −4.0 mm) or the DRN (AP = −5.5 mm, ML = ± 0 mm, DV = −2.0mm; at a 15° angle from caudal to rostral) for subsequent optogenetic activation and fiber photometry experiments, respectively. Post-surgery, the mice were maintained on a heating pad until they regained ambulatory function, after which they were returned to their home cages. Behavioral experiments were conducted three weeks later to allow for sufficient viral expression.

### Behavioral tests

#### Tube test

The tube test, a well-established methodology for evaluating social dominance among mice, has been extensively documented in previous research(Fan et al., 2019). In brief, following a co-housing period of at least two weeks in the same cage, four adult male C57BL/6 mice underwent three consecutive days of handling, succeeded by three consecutive days of tube test training. Training utilized an acrylic tube with an internal diameter of 3 cm and a length of 30 cm, with each mouse completing 10 tube entries per day (5 entries from each end). During the tube test, pairs of mice were placed at contrary ends of the tube, converging in the middle. The mouse that exited the tube first was designated as the loser, while its counterpart was labelled as the winner. Should neither mouse exit the tube within a two-minute period during a trial, the test was repeated. Each mouse engaged in three trials daily against its cage mates, culminating in a total of six trials per cage. The rankings of the four mice were subsequently determined based on the total number of victories achieved.

The behaviors of the animals during the tube test were meticulously captured on video and subsequently analyzed frame by frame using BORIS video analysis software (V7.12.2, University of Torino)(Friard and Gamba, 2016). In accordance with the behavioral classification criteria established in a previous study, the observed actions within the tube were categorized as follows: push initiation (actively instigating a push), push back (responding with a push after being challenged by an opponent), resistance (withstanding backward movement when confronted), stillness (exhibiting no movement beyond sniffing and grooming), and retreat (receding backward either in response to being pushed or by voluntary decision). Each distinct behavioral epoch was carefully annotated using BORIS to facilitate further analysis. To eliminate any potential bias in the analysis, the annotator was blinded to the hierarchical ranks or prior experiences of the mice during the behavioral annotation process.

#### Warm spot test

An adequate volume of water was poured into a rectangular plastic container (28 cm × 20 cm × 20 cm), which was subsequently placed in a freezer. Prior to the initiation of the experiment, a plastic film was affixed to the surface of the ice to maintain the temperature at approximately 0 ℃ throughout the duration of the test. A heating coil, encased in cardboard, was positioned in one corner of the container to elevate the local temperature to 34℃. This configuration was designed to simulate a warm nest with a diameter of 5 cm, accommodating a single adult mouse only. For the preparatory phase, four mice from a single cage were initially housed in a container with a base layer of ice, devoid of a warm nest, for 20 minutes to facilitate a reduction in their body temperature. Following this acclimatization period, the mice were transferred to the experimental container, equipped with both ice and a warm nest. The competition among the four mice for access to the warm nest was subsequently observed and recorded over a duration of 20 minutes.

The warm spot test was conducted on both hM4Di-EGFP mice and EYFP mice. One day following the initial test, the selected individual received an intraperitoneal injection of 10 mg/kg CNO, while its cage mates were administered an equivalent volume of saline solution. The warm spot test was subsequently repeated at 2 and 48 hours post-CNO injection. Using the BORIS software, we annotated the timestamps of entries into and evictions from the warm nest, and calculated the total duration each mouse occupied the nest. In instances where multiple animals occupied the warm nest simultaneously, only the mouse occupying the majority of the area within the warm nest was recorded. As with prior procedures, the annotator was blinded to the hierarchical ranks and past experiences of the mice during the behavioral annotation process.

#### Open field test (OFT)

To evaluate the motivation for exploring the central area of a novel environment and to monitor the locomotor activity of the test subjects, we gently placed each mouse in the center of a 40 × 40 cm open field apparatus and recorded its activity under dim light during the dark phase for a duration of 10 minutes. The open field test was conducted using a behavioral research system (CinePlex, Plexon, USA) for video recording and data acquisition. In optogenetic experiments, blue light (473 nm, 20 Hz, 20 ms, 10 mW) was intermittently activated for 1-minute epochs. For chemogenetic experiments, the open field test was performed 2 hours post-CNO injection. The tracking software (CinePlex Studio V3.5.0) was employed to calculate the number of entries into the central zone, the duration of stay in the central zone (center time), and the total distance traveled (locomotion) for each minute.

#### Elevated plus maze test (EPM)

Elevated plus maze assays were conducted to evaluate anxiety-like behaviors in mice utilizing a four-arm maze configuration comprising two open and two closed arms. Each mouse was placed in the central intersection of the maze, and their locomotor activities were recorded using the previously described system under low-light conditions during the dark phase over a 9-minute session. The duration spent in both open and closed arms was quantified using the CinePlex Studio software. In experimental groups, blue light (473 nm, 20 Hz, 20 ms, 10 mW) was intermittently activated for 3-minute intervals.

#### Social memory test

The social memory test was performed using a three-chamber apparatus (40 × 20 cm). Each test mouse was allowed to habituate in the central chamber for 10 minutes. Subsequently, an unfamiliar mouse was introduced into one of the side chambers. Following a 5-minute familiarization period with the introduced mouse, a novel object mouse was placed in the other side chamber. The subsequent 5-minute interval was used to observe the exploration behaviors of the test mice towards either the familiar or novel object mouse. For analysis, the time spent in the interaction zone (defined as half of the center chamber adjacent to the side chamber, 10 × 20 cm) was recorded in 1-minute epoch, during which the mice experienced alternating light-on and light-off periods (473 nm, 20 Hz, 20 ms, 10 mW, activated and deactivated intermittently in 1-minute epochs).

#### Resident-intruder test

The standard resident-intruder test was conducted on mice that had cohabitated with their cage mates for a duration exceeding one month. Prior to the assay, all cage mates were removed from the home cages, allowing the test mice to habituate for 10 minutes. Subsequently, adolescent male intruders (5 weeks old) were introduced into the cages for a 10-minute testing period. Concurrently, blue light (473 nm, 20 Hz, 20 ms, 10 mW) was intermittently administered in 1-minute epochs on the test mice. The frequency and duration of aggressive behaviors (e.g. chasing and attacking) and non-aggressive social interactions (e.g. sniffing and social grooming) exhibited by the test mice were recorded and analyzed for each 1-minute epoch corresponding to the light-on and light-off intervals.

#### Grip-strength measurement

We employed a force gauge (DS2-50N, Puyan, Dongguan, China) to measure the forelimb grip strength of mice. Each mouse underwent three testing blocks with 2-hour intervals between blocks. Within each block, mice completed 10 trials over approximately 6 minutes, comprising two equal 3-minute periods corresponding to light-on or light-off conditions (473 nm, 20 Hz, 20 ms, 10 mW). The sequence of light-on and light-off periods was varied pseudorandomly across the three blocks to mitigate potential order biases. Ultimately, the maximal grip strengths recorded from the three blocks under both light-on and light-off conditions were used for statistical analysis.

### Optogenetic manipulation of SNr and its downstream targets in the tube test

Following the establishment and maintenance of a stable rank in the tube test for over three days, all four mice were fitted with simulated optic fibers and underwent preliminary tube tests (featuring a 12 mm slit at the top) for two consecutive days prior to the main experiments. Photostimulation for each test subject commenced at 5 mW, with the tube test performed against the cage mate closest in rank, subsequently progressing to opponents with increasingly greater rank disparities. Mice were required to win or lose in two out of three trials from either end of the tube. If the test mice secured victories or experienced defeats from both ends of the tube, it was inferred that photostimulation at that specific intensity successfully induced a rank alteration. Conversely, if no rank change occurred, the light intensity was incrementally increased, and the procedure was repeated until a rank shift was observed or the light intensity reached 20 mW. The total number of winning trials induced by photostimulation in the test mouse was recorded. Subsequently, rank changes for each mouse were subjected to detailed statistical analyses, with rank 1 mice excluded due to the ceiling effect.

### Chemogenetic manipulation of SNr in the tube test

All four mice housed in the same cage were administered the rAAV-CaMKIIα-hM4D(Gi)-EGFP-WPRE-hGH-pA virus via injection. At least four weeks post-viral injection, the tube test was conducted at intervals of 1-1.5, 3-5, 6-8, 24, 48, and 72 hours following an intraperitoneal (i.p.) saline injection. One week later, the test mice received an i.p. injection of CNO (10 mg/kg), while the remaining mice were given saline injections. The tube test was then repeated at the specified time points. Test subjects were selected randomly. Following these procedures, rank changes for each mouse were evaluated through comprehensive statistical analyses, excluding rank 4 mice due to the floor effect.

### Fiber photometry recording and analysis

Ca^2+^ signals were recorded utilizing a fiber photometry system (QAXK-FPS-SS-LED, ThinkerTech, Nanjing, China). The GCaMP6f indicator was excited with 488 nm light, and the signals at 405 nm were subtracted from those at 488 nm to obtain corrected signals. Data were sampled at a frequency of 100 Hz, and the laser intensity was maintained at 40 μw at the tip of the optic fiber to minimize photobleaching. Behavioral observations during the tube test were captured using a camera positioned beside the tube and subsequently annotated with BORIS for further analysis. To synchronize the fiber photometry recordings with the video, an external trigger generated a TTL pulse to the recording system while simultaneously illuminating a red light-emitting diode, which was captured by the camera. Each pair of opponents underwent six trials, consisting of three entries from each side. As a control, mice were allowed to traverse the tube alone in six trials, with three entries from each side.

Ca^2+^ signals were analyzed using proprietary scripts provided by ThinkerTech Nanjing Biotech Co., Ltd., which were developed in MATLAB (2017b). Fluorescence changes, denoted as ΔF/F0, were calculated using the formula (F-F0)/F0, where F0 represents the baseline fluorescence averaged over a 2-second interval preceding the onset of a behavioral epoch. For peri-event time histogram (PETH) analysis, behavioral onset was aligned to time zero, and fluorescence signals were normalized using Z-score and binned at 10 ms intervals. Specifically, for the PETH analysis of mice traversing a tube, fluorescence signals were aligned to the moment that the mice reached the midpoint of the tube. Statistical significance of behavior-associated fluorescence changes was evaluated using a permutation test, following previously established methodologies.

### Histology and immunohistochemistry

Mice were anesthetized using 1% pentobarbital sodium and subsequently perfused transcardially, initially with saline and then with 4% paraformaldehyde. The brains were excised and fixed in 4% paraformaldehyde for a period of 24 hours, followed by storage in a 30% sucrose/phosphate buffer solution (PBS) at 4°C until sectioning. Coronal brain slices of 30 μm thickness were prepared for immunohistochemical staining using a cryostat (CM1850, Leica, Germany). Primary antibodies used in this study included rabbit anti-CaMKII α (1:300; Abcam, ab5683) and rabbit anti-TH (1:500, Millipore, AB152). Secondary antibody employed was donkey anti-rabbit Alexa Fluor 647 (1:500, Abcam, ab150111). The sections were subsequently mounted onto slides, coverslipped, and imaged using a fluorescence microscope (Nexcope NE910, Ningbo, China).

### Statistical analysis and Reproducibility

The normality of distributions and the homogeneity of variance for all data were assessed using the Shapiro-Wilk test and Levene’s test, respectively. For the tube test, the Mann-Whitney U test was employed to compare differences in behavioral parameters across various light conditions and among different groups. For the warm spot test, one-way repeated measures ANOVA with Bonferroni correction was used to compare the durations of occupying the warm spot, while the Friedman test was applied to compare ranks across different time points. For the open field test, one-way repeated measures ANOVA with Bonferroni correction was utilized to compare differences in behavioral parameters among groups subjected to chemogenetic manipulation, and paired-samples t-tests were conducted to compare behavioral parameters across different light conditions for optogenetic manipulation. For the elevated plus-maze and resident-intruder tests, two-way repeated measures ANOVA with Bonferroni correction was employed, with the variables being “light condition” and “group”. For the social memory test, two-way repeated measures ANOVA with Bonferroni correction was used, considering the variables “zone” and “group”. Paired-samples t-tests were conducted to compare forelimb grip strength between different light conditions. Statistical analyses were performed using SPSS software (release 23.0) with a significance threshold set at *p* < 0.05 and a highly significant threshold at *p* < 0.001. The reproducibility of micrographic and behavioral experiments in optogenetics, chemogenetics, and fiber photometry recordings was consistent with the actual number of experimental subjects, and the repeatability of micrograph statistics corresponded to the sample size. Furthermore, the repeatability of anterograde tracing experiments was confirmed to be at least three times.

## Data availability

The dataset generated and analyzed during the current study are available in the Mendeley repository: https://data.mendeley.com/drafts/xz92psvgbk

## Acknowledgements

This study was supported by the grants from the National Natural Science Foundation of China (No. 32170504).

## Author contributions

Y.-Z.F. and G.-Z.F. contributed the design of the experiments. Y.-Z.F. performed the surgery. S.-X.G., Y.-Z.F. and L.-D.L. performed the behavioral tests. S.-X.G. and Y.-Z.F. collected and analyzed the data. S.-X.G. and Y.-Z.F. performed histology and immunohistochemistry. W.-J.N., and Z.-Y.W., performed mice husbandry. Y.-Z.F. and G.-Z.F. wrote the manuscript. Y.-Z.F., X.-G.J. and G.-Z.F. contributed the textual revision of the paper. All authors discussed the paper.

