## Supplemental figures for "The Glutamatergic Projection from the Substantia Nigra Pars Reticulata to the Dorsal Raphe Nucleus Facilitates Social Hierarchy in Mice"

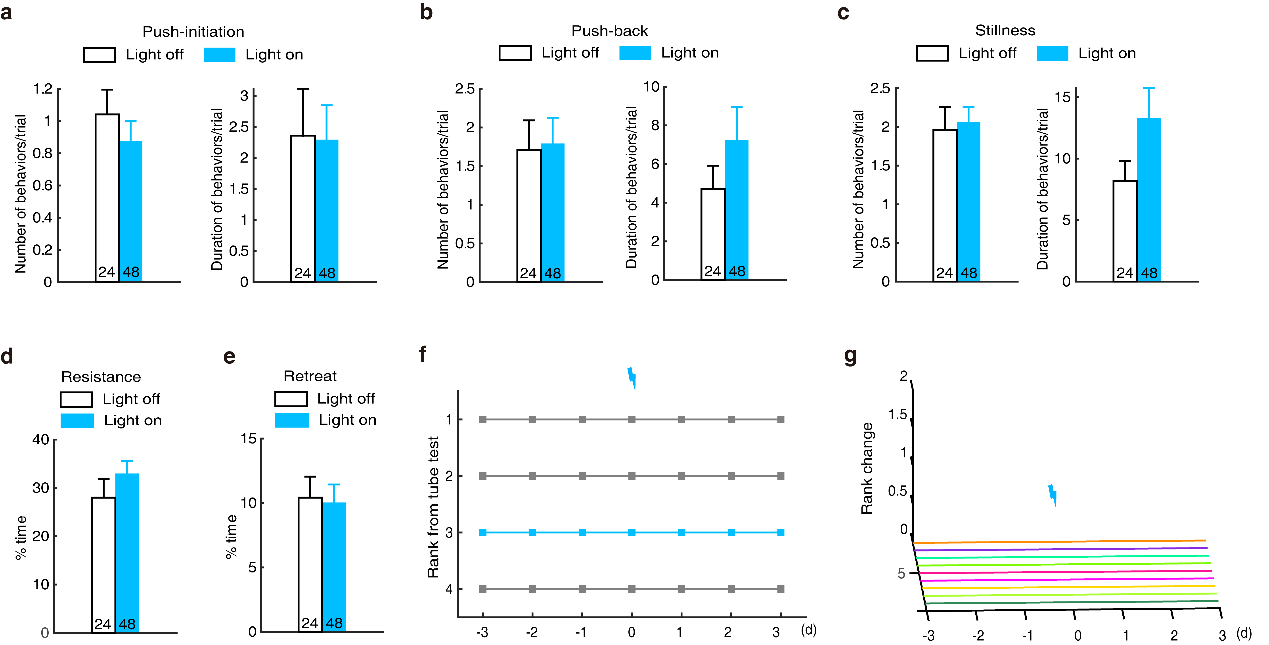

**Supplementary Figure 1. Photostimulation of SNr^Glu^ neurons did not affect behaviors and ranks of EYFP mice in the tube test.** **a** Number and duration of push-initiation for each trial in the tube test (n=24 for light off and n=48 for light on, Mann-Whitney U test; number: U=493.5, *p*=0.285; duration: U=487, *p*=0.280). **b** Number and duration of push-back for each trial in the tube test (n=24 vs. 48, Mann-Whitney U test; number: U=567.5, *p*=0.917; duration: U=556, *p*=0.808). **c** Number and duration of stillness for each trial in the tube test (n=24 vs. 48, Mann-Whitney U test; number: U=543.5, *p*=0.689; duration: U=478.5, *p*=0.244). **d** Percentage of time spent resisting (n=24 vs. 48, Mann-Whitney U test; U=490, *p*=0.304). **e** Percentage of time spent retreating (n=24 vs. 48, Mann-Whitney U test; U=528, *p*=0.566). **f** Example of rank positions for one cage of mice tested daily over 7 days, showing that the third-ranked mouse did not change its rank following photostimulation of SNr^Glu^ neurons. **g** Summary of rank changes in EYFP mice before and after photostimulation on SNr^Glu^ neurons (n=9). Each line represents one animal.

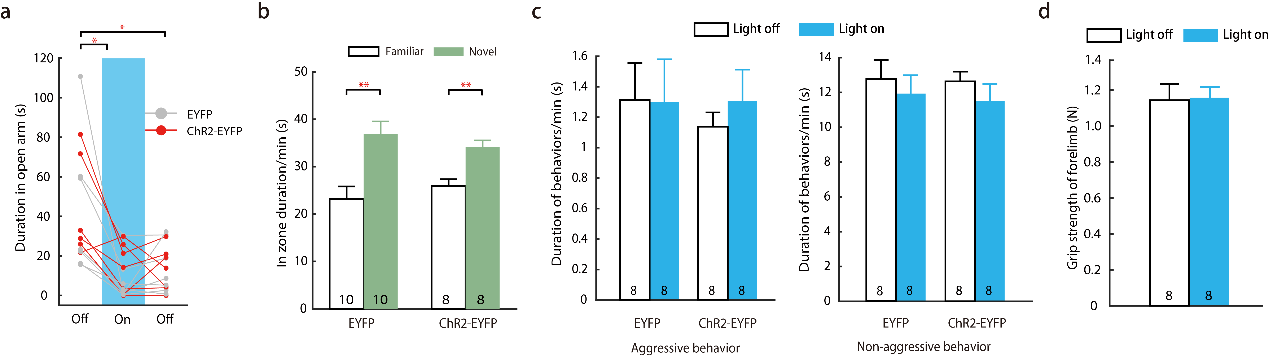

**Supplementary Figure 2. Optogenetic activation of SNr^Glu^ neurons did not impact anxiety levels, social preference, aggressive behavior, and grip strength of mice. a** During the elevated plus maze (EPM) test, time spent in the open arm decreased over time (n=6 for ChR2-EYFP mice and n=7 for EYFP mice, two-way repeated measures ANOVA with Bonferroni correction; F_2,22_=15.814, *p*<0.001), though no significant differences were observed between groups (F_1,11_=0.621, *p*=0.447). **b** In the social memory test, both ChR2-EYFP and EYFP mice showed a significant preference for novel mice during light-on period (n=8 and 10 for ChR2-EYFP and EYFP mice respectively; two-way repeated measures ANOVA with Bonferroni correction; F_1,16_=11.293, p=0.004). However, no significant differences were found between groups (F_1,16_=4.127, *p*=0.059). **c** The resident-intruder test indicated that both ChR2-EYFP and EYFP mice displayed normal levels of aggressive and non-aggressive behaviors towards novel mice (n=8 for each group, two-way repeated measures ANOVA with Bonferroni correction; F_1,14_=0.146, *p*=0.708), with no significant differences between groups (F_1,14_=0.137, p=0.716). **d** Optogenetic activation of SNr^Glu^ neurons did not affect forelimb grip strength during light-on and light-off periods (n=8, paired two-sided t-test; t7=1.735, *p*=0.126). Data represented as mean ± SEM. **p* < 0.05; ***p* < 0.01.

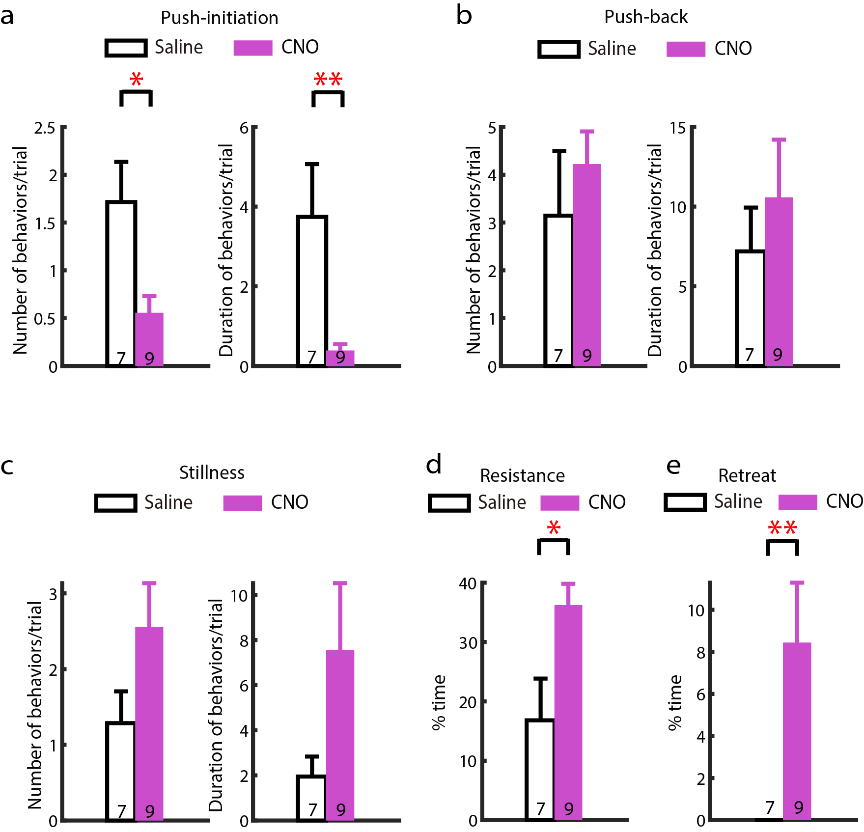

**Supplementary Figure 3. Chemogenetic inhibition of SNr^Glu^ neurons in hM4Di-EGFP mice reduces effortful behaviors in the tube test 3.5 hours post-CNO injection compared to saline injection. a** Number and duration of push-initiation for each trial in the tube test (n=7 mice for saline injection and n=9 mice for CNO injection, Mann-Whitney U test; number: U=10, *p*=0.011; duration: U=5, *p*=0.005). **b** Number and duration of push-back for each trial in the tube test (n=7 vs. 9, Mann-Whitney U test; number: U=20, *p*=0.218; duration: U=26, *p*=0.560). **c** Number and duration of stillness for each trial in the tube test (n=7 vs. 9, Mann-Whitney U test; number: U=18, *p*=0.140; duration: U=13, *p*=0.050). **d** Percentage of time spent resisting (n=7 vs. 9, Mann-Whitney U test; U=12, *p*=0.039). **e** Percentage of time spent retreating (n=7 vs. 9, Mann-Whitney U test; U=7, *p*=0.004). Note that the same 3 animals were used in both groups; specifically, 3 out of 9 mice received i.p. injection of saline one week prior to the CNO injection. Data represented as mean ± SEM. * *p* < 0.05; ** *p* < 0.01.

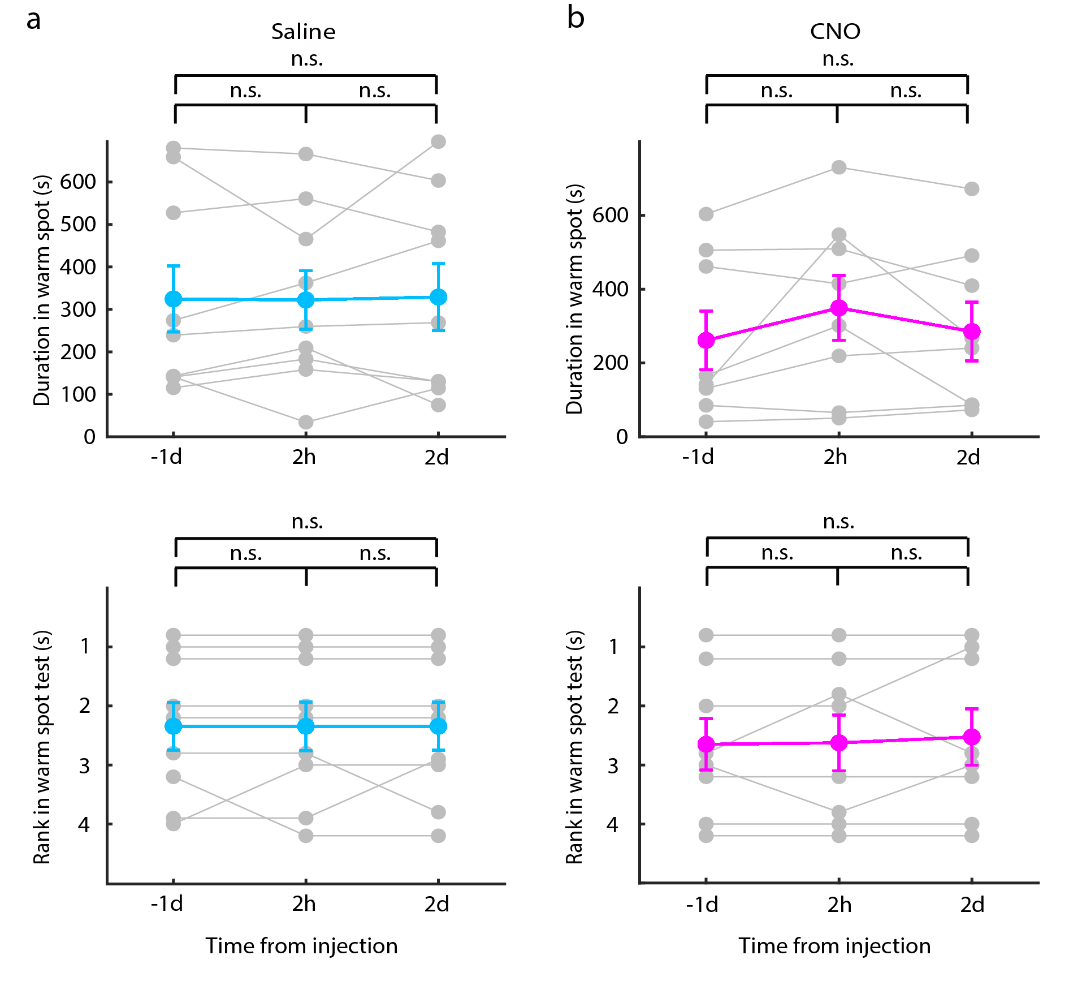

**Supplementary Figure 4.** **Chemogenetic manipulation of SNr^Glu^ neurons in EYFP mice did not influence the duration of occupation or ranking in the warm spot test following injections of either saline or CNO.** **a** The duration (top) and rank (bottom) in the warm spot test were assessed for mice expressing EYFP in the SNr at 1 day before, 2 hours post, and 2 days post i.p. administration of saline (duration in the warm spot: n=9, one-way repeated measures ANOVA with Bonferroni correction, F_2,16_=0.007; *p*=0.993; rank: Friedman test, t_2_=0.000; *p*=1.000). **b** Similarly, the duration (top) and rank (bottom) in the warm spot test for mice expressing EYFP in the SNr at 1 day before, 2 hours after, and 2 days following i.p. injection of CNO (duration: n=8, one-way repeated measures ANOVA with Bonferroni correction, F_2,14_=2.273; *p*=0.140; rank: Friedman test, t_2_=0.667; *p*=0.717). Notably, all mice received an i.p. injection of saline one week prior to the CNO injection. Each line on the graph represents an individual animal.

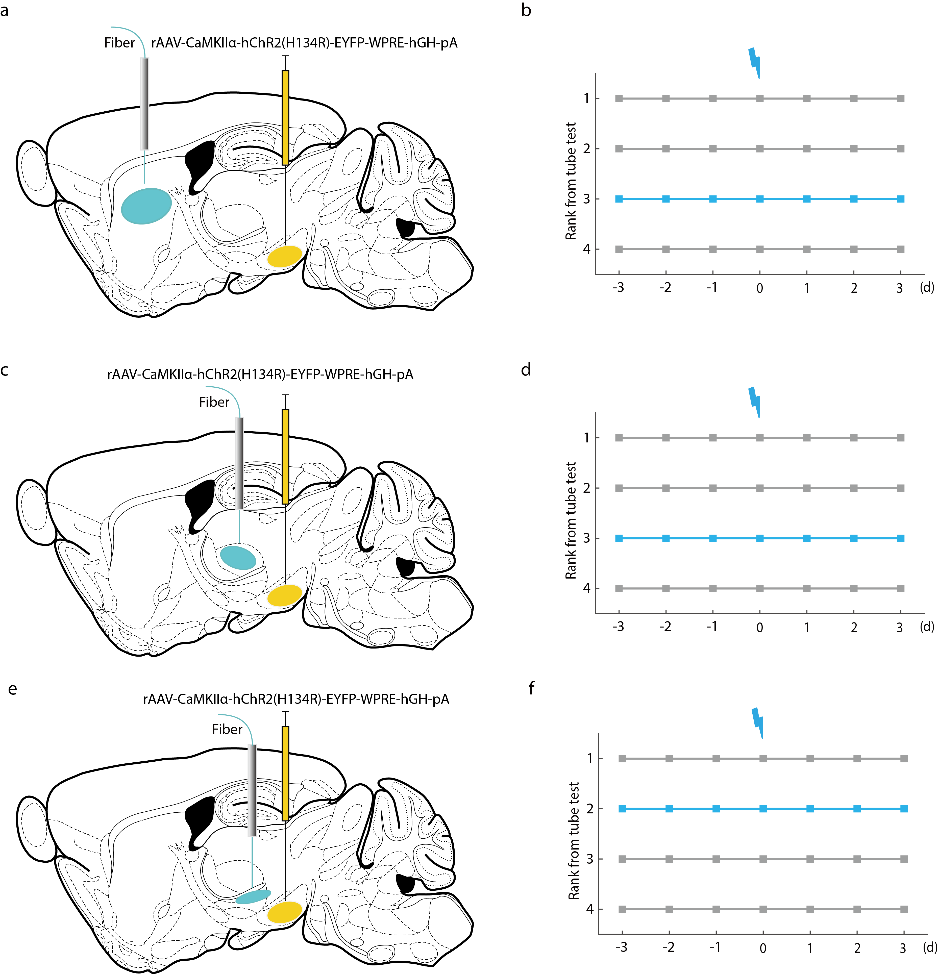

**Supplementary Figure 5.** **The optogenetic stimulation of the SNr^Glu^-CPu, -VPM and -ZID pathways did not influence the social hierarchy of mice.** Panels **a, c** and **e** illustrate diagrams of the unilateral viral infection area using rAAV-CaMKIIα-hChR2(H134R)-EYFP-WPRE-hGH-pA in the SNr, along with the placement of optic fibers above the CPu (a), VPM (c) and ZID (e), respectively. Panels **b, d** and **f** present examples of rank positions for a group of mice assessed daily over seven days, showing that the rank of the third-position mouse remained unchanged following photostimulation of SNr^Glu^ neuronal terminals in the CPu (b), VPM (d) and ZID (f), respectively. Each line represents an individual animal. Abbreviations: CPu, caudate putamen (striatum); DRN, dorsal raphe nucleus; VPM, ventral posteromedial thalamic nucleus; ZID, zona incerta, dorsal part. The stereotaxic coordinates (in mm) are: CPu (AP = 0.5, ML = ± 2.0, DV = -3.0), VPM (AP = -1.0, ML = ± 0.8, DV = -3.5) and ZID (AP = -1.8, ML = ± 1.0, DV = -4.0), DRN (AP = -5.5, ML = ± 0, DV = -2.50).

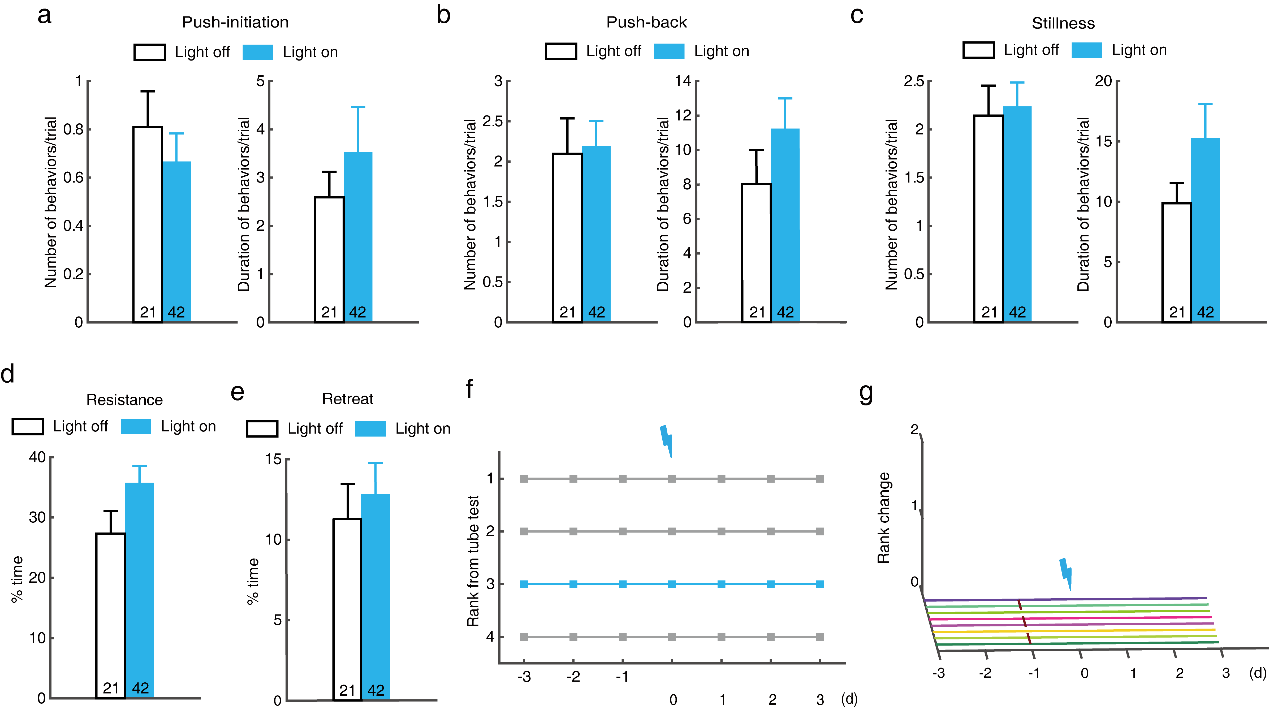

**Supplementary Figure 6. Photostimulation of the SNr^Glu^-DRN pathway did not significantly alter the behaviors or hierarchical ranks of EYFP mice. a** Number and duration of push-initiation for each trial in the tube test (n=21 for light off and n=42 for light on, Mann-Whitney U test; number: U=381.5, *p*=0.342; duration: U=391.5, *p*=0.452). **b** Number and duration of push-back for each trial in the tube test (n=21 vs. 42, Mann-Whitney U test; number: U=416.5, *p*=0.715; duration: U=361.5, *p*=0.245). **c** Number and duration of stillness for each trial in the tube test (n=21 vs. 42, Mann-Whitney U test; number: U=438, *p*=0.964; duration: U=414, *p*=0.694). **d** Percentage of time spent resisting (n=21 vs. 42, Mann-Whitney U test; U=329, *p*=0.102). **e** Percentage of time spent retreating (n=21 vs. 42, Mann-Whitney U test; U=427, *p*=0.838). **f** Daily rank positions of a sample cage of mice over 7 days, illustrating that the third-ranked mouse maintained its rank following photostimulation of SNr^Glu^ neuron terminals in the DRN. **g** No changes in rank were observed in EYFP mice before and after the optogenetic activation of the SNr^Glu^-DRN pathway. Each line represents an individual animal. Data represented as mean ± SEM.

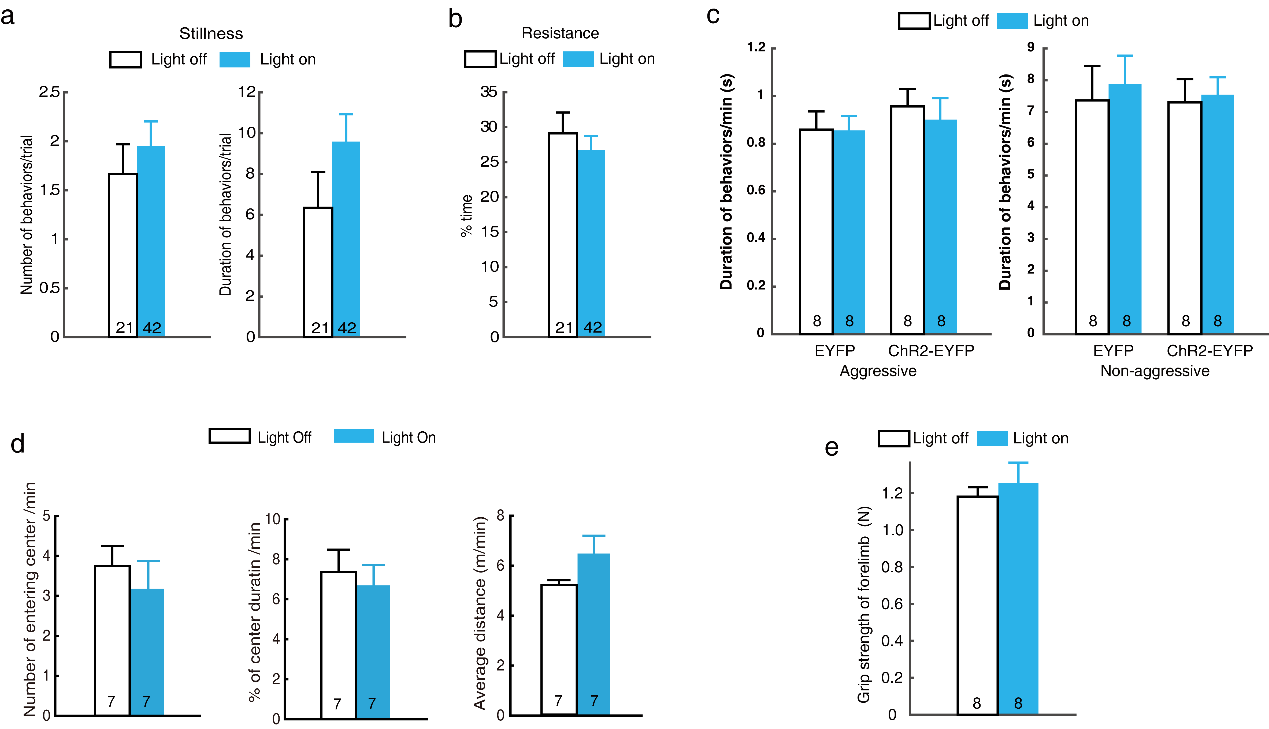

**Supplementary Figure 7. Photostimulation of the SNr^Glu^-DRN pathway did not influence other behavioral parameters in ChR2-EYFP mice. a** Number and duration of stillness for each trial in the tube test for ChR2-EYFP mice (n=21 for light off and n=42 for light on, Mann-Whitney U test; number: U=406, *p*=0.596; duration: U=331, *p*=0.108). **b** Percentage of time spent resisting (n=21 vs. 42, Mann-Whitney U test; U=389, *p*=0.448). **c** Both ChR2-EYFP and EYFP mice demonstrated typical levels of aggressive (n=8 for each group, two-way repeated measures ANOVA with Bonferroni correction; F_1,14_=0.173, *p*=0.684) and non-aggressive behaviors (F_1,14_=0.207, *p*=0.656) in the resident-intruder test, with no significant differences observed between the groups (F_1,14_=0.782, *p*=0.391 for aggressive behaviors; F_1,14_=0.06, *p*=0.811 for non-aggressive behaviors). **d** Optogenetic activation of the SNr^Glu^-DRN pathway did not alter the time spent in the center (n=7, paired two-sided t-test, t_6_=0.941, *p*=0.383), overall locomotion (n=7, Wilcoxon signed rank test, Z_6_=-1.521, *p*=0.128), or the number of entries into the central area (n=7, paired two-sided t-test, t_6_=1.307, *p*=0.239) during the open field test. **e** Optogenetic activation of the SNr^Glu^-DRN pathway did not alter forelimb muscle strength during both light-on and light-off periods (n=8, paired two-sided t-test; t_7_=0.643, *p*=0.541). Data represented as mean ± SEM.
